# Electrophysiological correlates of state transition prediction errors

**DOI:** 10.1101/544551

**Authors:** Danesh Shahnazian, José J.F Ribas-Fernandes, Clay B. Holroyd

## Abstract

Planning behavior depends crucially on the ability to distinguish between the likely and unlikely consequences of an action. Formal computational models of planning postulate the existence of a neural mechanism that tracks the transition model of the environment, i.e., a model that explicitly represents the probabilities of action consequences. However, empirical findings relating to such a mechanism are scarce. Here we report the results of two electroencephalographic experiments examining the neural correlates of transition model learning. The results implicate fronto-midline theta and delta oscillations in this process and suggest a role of the anterior midcingulate cortex in planning behavior.

## 1. Introduction

Planning is a form of decision making that involves simulating possible outcomes of potential actions and selecting the action that is most likely to result in a desired outcome (also referred as goal-directed decision making and model-based reinforcement learning; Daw, 2018). Planning is a more flexible mode of decision making in human and nonhuman animals, which allows the consideration of rarely experienced actions and outcomes (Daw, Gershman, Seymour, Dayan, & Dolan, 2011), or by allowing a rapid adaptation to a change in action-outcome contingencies (Dickinson and Balleine, 2002). This stands in contrast with stimulus-response learning (or model-free reinforcement learning), where an agent learning stimulus-response associations often perseverates on previously rewarded but currently suboptimal behaviors (Daw, Niv & Dayan, 2005). Decades of research on the neural mechanisms of planning have consistently implicated the dorsolateral prefrontal cortex (DLPFC) and the hippocampus in some aspects of this complex behavior (Unterrainer, & Owen, 2006; Foster and Wilson, 2006; Johnson and Redish, 2007; Smittenar, 2013), but other questions have yet to be addressed. For example, only a few studies have focused on how the brain represents multiple possible outcomes of a single action (e.g, Hamilton & Grafton, 2007). The complexity of planning strongly suggests that this cognitive process is supported by a complex network of brain areas.

Notably, recent research has associated the anterior midcingulate cortex (aMCC) with planning. In a seminal study, Balaguer, Spiers, Hassabis and Summerfield (2016) recorded the fMRI BOLD response from participants engaged in a hierarchical planning task that required navigating through a simulated subway network. On each trial, participants started at a randomly selected station in the network and were tasked with finding a specific goal station, with each trial offering 4 potential actions corresponding to the cardinal directions. They observed that the BOLD activity in the aMCC and the premotor cortex increased linearly with the complexity of the plan that was being executed. This is consistent with a study by Simon and Daw (2011) that reported a similar increase in activity in both the DLPFC and the aMCC during a spatial navigation task.

Crucially, aMCC may play a role in planning by predicting the outcomes of the actions currently being taken, including their probability of occurrence, namely whether the outcomes are common or rare. One way to represent such associations is through constructing a *transition model*. A transition model reflects the agent’s expectations about the probability that an upcoming state of the environment is associated with the currently executed action in the context of the current state (Daw, Niv, & Dayan, 2005). Algorithms that use such explicit transition models are called model-based algorithms. This is in contrast to the well-known family of model-free algorithms (e.g., temporal difference learning), which are only concerned with associating actions with the expected future reward (Sutton & Barto, 1998). Formal models of planning rely on an explicit representation of the transition model. The transition model is updated based on state transition errors (SPEs) (Gläscher et al, 2010), which reflect discrepancies between predicted and actual outcomes and are larger for state transitions that are less expected relative to those that are more expected.

Gläscher et al. (2010) conducted an fMRI experiment using a paradigm that featured common and uncommon transitions and found that the intraparietal sulcus and lateral PFC are sensitive to SPEs. However, this result seems to contrast with the predictions of Alexander and Brown’s Predicted Response-Outcome (PRO) model (2011), which casts the aMCC as a predictive mechanism that signals violations of predicted action-outcome contingencies. The model has successfully accounted for a variety of empirical results obtained using different methodologies ranging from single cell studies to fMRI studies. The model represents all the possible outcomes of each potential action explicitly and estimates the likelihood of each of these outcomes. When the specific outcome of an action is observed, the likelihood estimates are updated accordingly. According to the model, the univariate recordings from the aMCC reflect the sum of these update signals. The model therefore designates the aMCC as a domain-general mechanism that generates prediction errors. These prediction errors, which inform different action-selection mechanisms, are very similar to the SPEs discussed earlier. Therefore, the notion that the aMCC is involved in learning a transition model to inform planning is compatible with this model (see also Shahnazian & Holroyd, 2018; Vassena, Holroyd & Alexander, 2018).

Consistent with these theories, several human functional neuroimaging (e.g., O’Reilly et al., 2013) and electrophysiological (e.g., Wessel et al., 2012) phenomena associated with aMCC have been shown to reflect discrepancies between predicted and actual events. Of note, fronto-midline theta (FMT) oscillations in the electroencephalogram (EEG), which appears to be generated in the aMCC (Ishii et al., 1999), are elevated to infrequently occurring novel, oddball (Cavanagh, Zambrano Vazquez & Allen, 2012) and feedback (HajiHosseini & Holroyd, 2012) stimuli. The sensitivity of FMT to event probability suggests that this signal could reflect the detection of SPEs by aMCC.

Whereas only a few studies have focused on the neural implementation of transition models, several studies have examined how planning algorithms modulate action-value representation in the brain (for a review, see Doll and Dayan, 2013). Planning algorithms work essentially by computing the expected value of each possible course of action, i.e., the sum of the transition probabilities associated with each possible consequence of an action multiplied by their predicted reward value. In this way, the planning algorithm evaluates the value of each action by estimating the expected reward associated with each action from the observed reward on each episode. The transition model modulates the value representations for different actions. Some theories have postulated that the brain utilizes a model-based reward prediction signal that computes the discrepancy between the expected reward based on incorporating the transition model and the actual reward that is observed in the current episode.

Inspired by this idea, Daw et al (2011) employed the *two-step choice task* in an fMRI experiment to tease apart model-based influences on participants’ choice behavior and its associated brain activity. Specifically, they hypothesized that model-based value should modulate a model-based reward prediction error (RPE) signal: Because an explicit model of the state-transitions changes the agent’s expectation, such an RPE should be modulated by these expectations. Yet contrary to their predictions, this RPE signal was localized to ventral striatum rather than to medial prefrontal cortex. Therefore, the exact contribution of the medial prefrontal cortex—and aMCC in particular—to model-based planning remains unclear.

We therefore set out to investigate this question by examining the time-frequency correlates of SPEs in the ongoing EEG. Here we present the results of two EEG experiments based on variants of the two-steps choice task. Although the two-step choice task provides several advantages in eliciting planning behavior in laboratory settings, it also poses a specific challenge for studying of the neural correlates of state transitions: the state transitions can simultaneously elicit both SPE and RPEs, especially when the participants select the first of the two sequential actions on each trial. Assuming that they expect the probable outcome of that first action to be more valuable than the rare outcome, the actual occurrence of the rare outcome should elicit both a negative RPE (given that it is undesired) and an elevated SPE (given that it is unpredicted). Thus, a strong discrepancy signal elicited by such events could reflect a SPE, an RPE, or both.

For this reason, in Experiment 1 we simplified the paradigm so that it elicits only SPEs (and not RPEs). We predicted that FMT power would be enhanced to transitions that are less frequent relative to the ones that are more frequent, thus highlighting the role of aMCC in the processing of SPEs. Then, in Experiment 2, we conducted the two-step choice task itself (Daw et al., 2011), in order to determine the degree to which the results of Experiment 1 would generalize to the more standard version of the task. Note that Experiment 1 does not incorporate rewards so is not a study of planning per se, whereas Experiment 2 does investigate planning.

## 2. Experiment 1

### 2.1. Method

#### 2.1.1. Participants

20 participants (12 females; 19 right handed; aged 17–30, *M =* 21.2, *SD* = 2.8) participated in the experiment. All of the participants had normal or corrected-to-normal vision and none reported a history of head injury. Participants were undergraduate students recruited from the University of Victoria. Each received course credit as well as a monetary bonus based on their task performance, as described below. All participants gave informed consent. The study was approved by the local research ethics committee and was conducted in accordance with the ethical standards prescribed in the 1964 Declaration of Helsinki.

#### 2.1.2. Task design

The computerized task was comprised of 6 blocks of 108 trials. On each trial, participants selected between images of 3 spaceships with distinct colours by pressing a key on a standard Qwerty keyboard (self-paced 2 second time-out) (Figure 1). Each spaceship was 3 cm wide and 6 cm long. Participants were seated 72 cm away from the screen and the center of the spaceship was at a distance of 13 cm from the center (visual angle 10 degrees). The stimuli presented to the participants included the image of two planets. Each planet-image’s radius on the screen was 16 cm (Note that the experimental code and visual stimuli for this task was adapted from alien two-step task with permission, see Decker, Otto, Daw, & Hartley (2016)).

**Figure 1.**
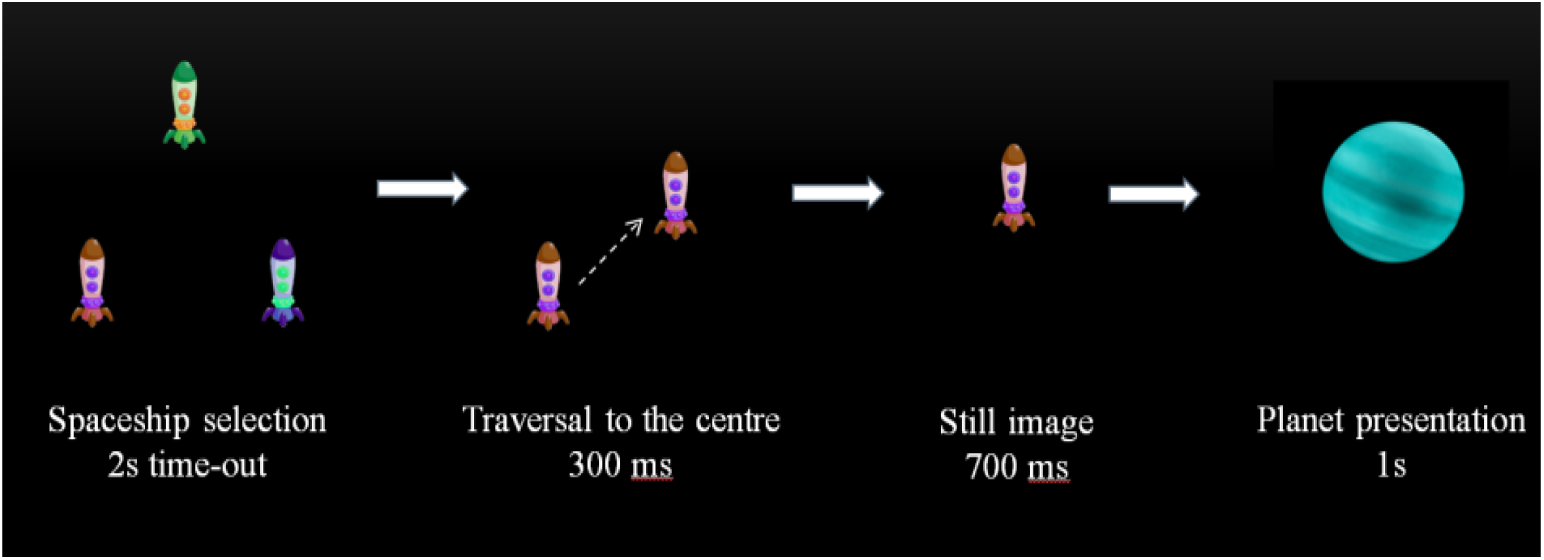
Sequence of events for an example trial. Participants selected between 3 spaceships spaced evenly on the vertices of an imaginary triangle. The selected spaceship then traversed to the centre of the screen (300 ms), stayed there (700 ms), and was then replaced by a planet image (1 s). White arrows indicate the event sequence. The dashed arrow indicates movement of a selected spaceship to the centre of the screen. The time presented underneath each event indicates its duration.

The spaceships were presented on the corners of an equilateral triangle with its centroid located at the centre of the screen. Participants pressed keys “h”, “j”, and “k” to select the spaceships on the bottom left, top, and bottom right corners of the triangle, respectively. They were instructed to rotate their selection through the 3 spaceship choices in a clock-wise order across trials. After a selection was made, the unselected spaceship disappeared and the selected spaceship moved from its location at a corner of the triangle directly to the centre of the screen, which served to guide the participant’s gaze to that location (a transition of 300 ms). After another 700 ms, the selected spaceship disappeared and the image of either a gold-coloured planet or a cyan-coloured planet appeared on the screen at that location, which simulated “travel” to the planet. Participants were instructed to attend to the identities of the selected spaceship and planet to which it travelled (Figure 1).

Crucially, on each block of trials, two of the three spaceships were more frequently associated with one of the two planets relative to the other planet. Specifically, selecting one spaceship was more likely (66%) to result in the appearance of the image of the gold planet while selecting one of the other two spaceships was more likely (66%) to result in the appearance of the image of the cyan planet. The third, remaining spaceship was associated with both planets with equal probability. The spaceship-planet mappings were pseudo-randomly determined across blocks, and never repeated across consecutive blocks. Therefore, participants learned such associations anew on each block of trials.

To ensure that participants attended to the associations between the spaceships and planets, they were probed at the end of each block for a total of six probes. Participants were presented with the image of one planet and were then presented with the image of the three spaceships. They were required to select the corresponding spaceship using the same response mappings as during the block. Participants were rewarded a $1 CAD bonus per correct answer, which was paid to them at the end of the experiment. The identity of the probe planet was pseudo-randomly determined for each block.

#### 2.1.3. Analysis of behavioral data

To investigate whether participants paid attention to the association between the selected spaceship and the upcoming planet, we analyzed the participants’ ratios of correct choices to all choices on the test probe. Chance performance on average would result in 1/3 correct responses; therefore, the mean for the null hypothesis distribution for accuracy is around 0.33. We assumed that the null-distribution is a t-student distribution.

#### 2.1.4. EEG acquisition and preprocessing

The EEG signal was acquired using a montage of 33 Ag/AgCl ring electrodes mounted on a nylon electrode cap according to the extended international 10–20 system (Jasper, 1958). Inter-electrode impedances were maintained below 20 kΩ using an abrasive conductive gel applied to each electrode. The electro-occulogram (EOG) was recorded for the purpose of ocular correction; horizontal EOG was recorded from the external canthi of both eyes, and vertical EOG was recorded from the sub-orbit of the right eye and electrode channel Fp2. Signals were amplified by differential amplifiers with a frequency response of DC 0.017–67.5 (90 dB per octave roll off) and digitized with a sampling rate of 250 per second. Digitized signals were stored on disk using Brain Vision Recorder software (Brain Products GmbH, Munich). Post-processing and data visualization were performed using Brain Vision Analyzer software (Brain Products GmbH, Munich). A fourth order digital Butterworth passband filter in the range of 0.1–20 Hz was applied (12 dB per octave roll off). Ocular artifacts were corrected using the eye movement correction algorithm described by Gratton, Coles, and Donchin (1983).

#### 2.1.5. Time frequency analysis

To estimate the time-frequency responses to the transition stimuli from EEG data, 2.4-s epochs extending from 1.4 s before to 1 s after stimulus onset (common and rare transition stimuli) were extracted from the single-trial data and convolved with a complex Morlet wavelet using BrainAnalyzer:

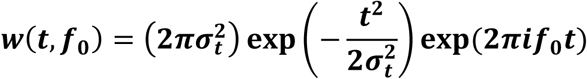

In this equation, t denotes time since the onset of the epoch, *f*_0_ denotes frequency of interest, 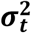 the variance of the normal distribution and *i* the imaginary unit. The wavelet family (2*πσ*_*t*_*f*_0_) ratio was set to 4. Changes in power spectrum over time (squared amplitude of the convolution between the signal and the wavelet) in 8 frequency points in the 2–10 Hz interval were computed for each single trial and averaged for each subject and condition before creating grand averages across subjects. To remove any possible baseline differences between conditions, the average baseline activity (300 ms–100 ms pre-stimulus) for each condition was subtracted from each data point following stimulus presentation for the corresponding frequency. The total difference in theta power elicited by transition stimuli for each subject was then calculated by averaging the amplitude of the difference in the time-frequency response to conditions (rare-common) in the 5–7 Hz frequency range and the 200-360 ms post-stimulus interval (HajiHosseini & Holroyd, 2012).

### 2.2. Results

#### 2.2.1. Behavioral data

Participants made the correct choice with a frequency that was significantly higher than chance level of performance (a paired samples t-test on the ratio of correct choices to all choices on the test probe revealed a significant difference between the sample mean and the mean for the sampling distribution assumed under the null hypothesis (*M* =.70, *SD* =.14), *t*(19) = 11.1, *p* < .05).

#### 2.2.2. Time-frequency response

Figure 2 shows the grand-average difference in the time-frequency response to the transition stimuli in the 2–10 Hz frequency range (top panel), and the scalp distribution of FMT power (bottom panel). As can be seen by inspection, theta power exhibits a fronto-central distribution, consistent with its identification as FMT.

**Figure 2.**
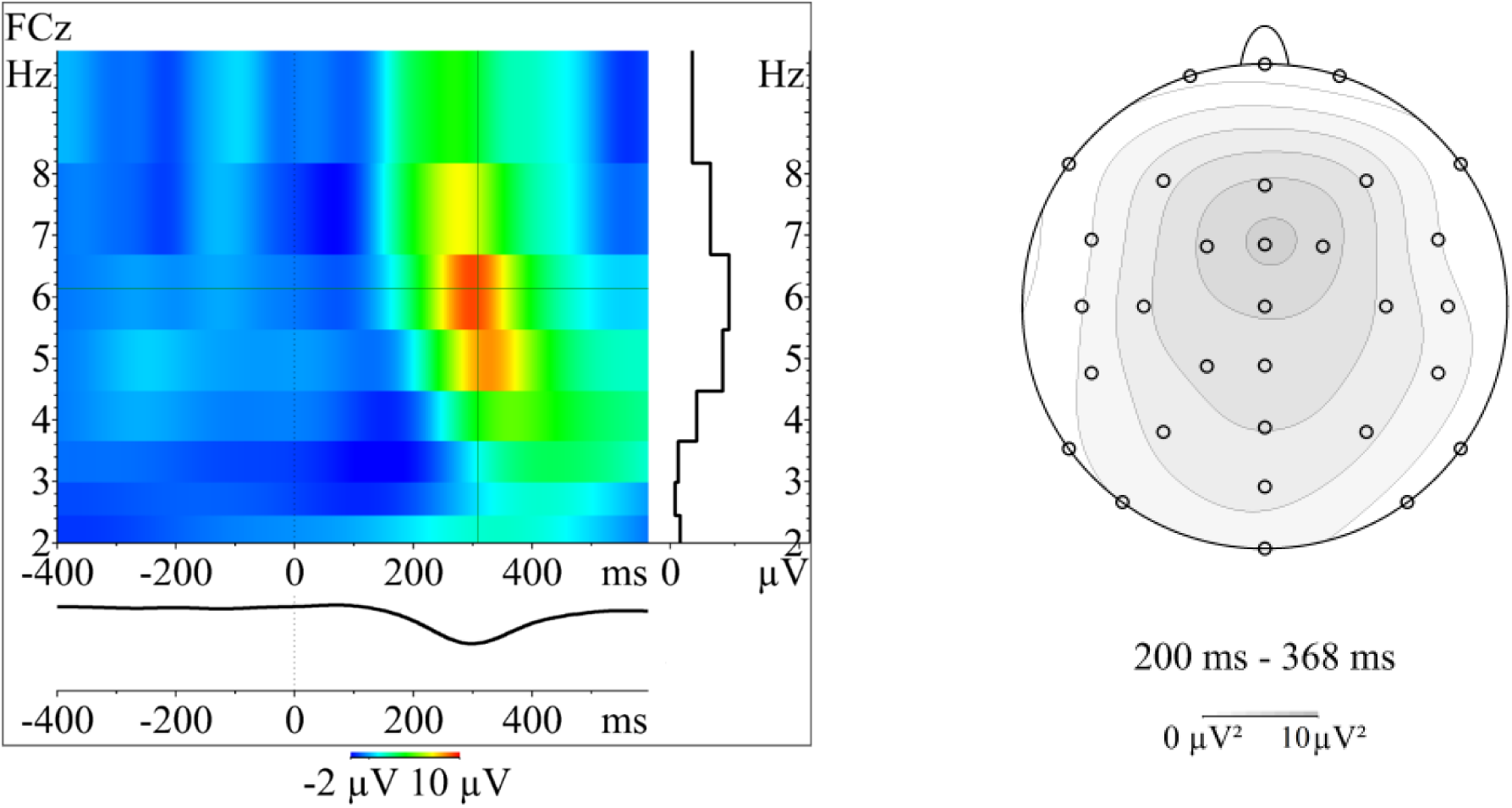
Left. The amplitude of difference in power in the time-frequency response to rare versus common transition (i.e. rare-transition) at different timepoints and frequencies in the range 2–8 Hz. Note that the crosshairs denote the timing and the frequency of maximum power. Right. The scalp distribution associated with the average difference in time-frequency response in the 4–7 Hz range 200-360 ms post-stimulus.

The mean difference in FMT power as a function of transition type (computed by subtracting the response to common transitions from that of rare transitions) was significantly different from zero (*M* = 7.47, *SD* = 2.34), *t*(19) = 6.44, *p* < .05).

Further exploratory analyses indicated that the amplitude of two ERP components (the N2 and the P3) were also larger to the rare transitions relative to the common transitions (see supplementary online material).

### 2.3. Interim Discussion

In this experiment, we used a variant of the two-step task where participants were instructed to track the likelihood of each upcoming state given the taken action. Participants’ choice behavior showed that they were successful at distinguishing between the likelihood of states given each action. Importantly, FMT power was enhanced by the unlikely outcomes relative to the likely outcomes following each action, implicating the signal as a product of transition prediction errors, as we predicted. Further, the scalp distribution of the theta signal, as isolated by the difference between the conditions, was fronto-central. This is consistent with the identification of this theta signal as FMT, as we predicted. These findings implicate the aMCC in the processing of the SPEs.

## 3. Experiment 2

Although we conducted Experiment1 to investigate the neural correlates of SPEs, the task did not allow for studying model-based planning because of the absence of reward (the object of planning). For this reason, in Experiment 2 we recorded EEG from participants engaged in a standard two-step choice task, in which the outcome of the first choice on each trial informs how a model-based agent should choose on the second step of the trial in order to obtain a reward. Therefore, participants who adopt a model-based strategy in this task should be sensitive to the likely and unlikely outcomes following the choice on the first step. We expected that the modulations of ERPs to the first step choice outcome in Experiment 2 should be similar to the pattern of results observed for Experiment 1, namely, greater FMT to rare relative to common state transitions.

That said, an important difference between Experiment 1 and 2 is that the RPE and SPE signals are concurrently modulated by the transition event. This could confound our analysis of the EEG data and possibly lead to discrepant results between the two experiments. Thus, Experiment 1 provided a meaningful control that can help disentangle the possible effects of transition errors in planning whereas Experiment 2 allowed us to evaluate this in the presence of dynamic reward learning.

### 3.1. Method

#### 3.1.1. Participants

40 participants (28 females; 31 right-handed; aged 17–31, *M* = 22.2, *SD* = 3.1) participated in the experiment. All of the participants had normal or corrected-to-normal vision and none reported a history of head injury. Participants were undergraduate students recruited from the University of Victoria. Each received course credit as well as a monetary bonus based on their task performance, as described below. All participants gave informed consent. The study was approved by the local research ethics committee and was conducted in accordance with the ethical standards prescribed in the 1964 Declaration of Helsinki.

#### 3.1.2. Task design

As mentioned in the introduction, the task is a modified version of the two-step task (Daw et al., 2011). According to the cover story delivered as task instruction to the participants, they are to travel to a planet (first-step choice) and select an alien from on that planet (second-step choice) to mine for treasure. Accordingly, the main difference between the two versions is that we used images of planets and aliens as stimuli (see below). The task consisted of 5 blocks of 50 trials (250 trials overall) with each trial involving two sequential decisions. According to the task instructions, the first decision involved choosing between two spaceship images. Subjects were told that these spaceships could visit either a “red” planet or a “purple” planet, with one spaceship visiting the red planet more often (70%) and the other one visiting the purple planet more often (70%). Figure 3 indicates the transition probabilities of each spaceship, which were explicitly instructed to the participants (in contrast to Experiment 1). The second decision involved choosing between two alien images on each planet (4 aliens in total). Subjects were told that the aliens dig for treasure and are differentially likely to be successful.

**Figure 3.**
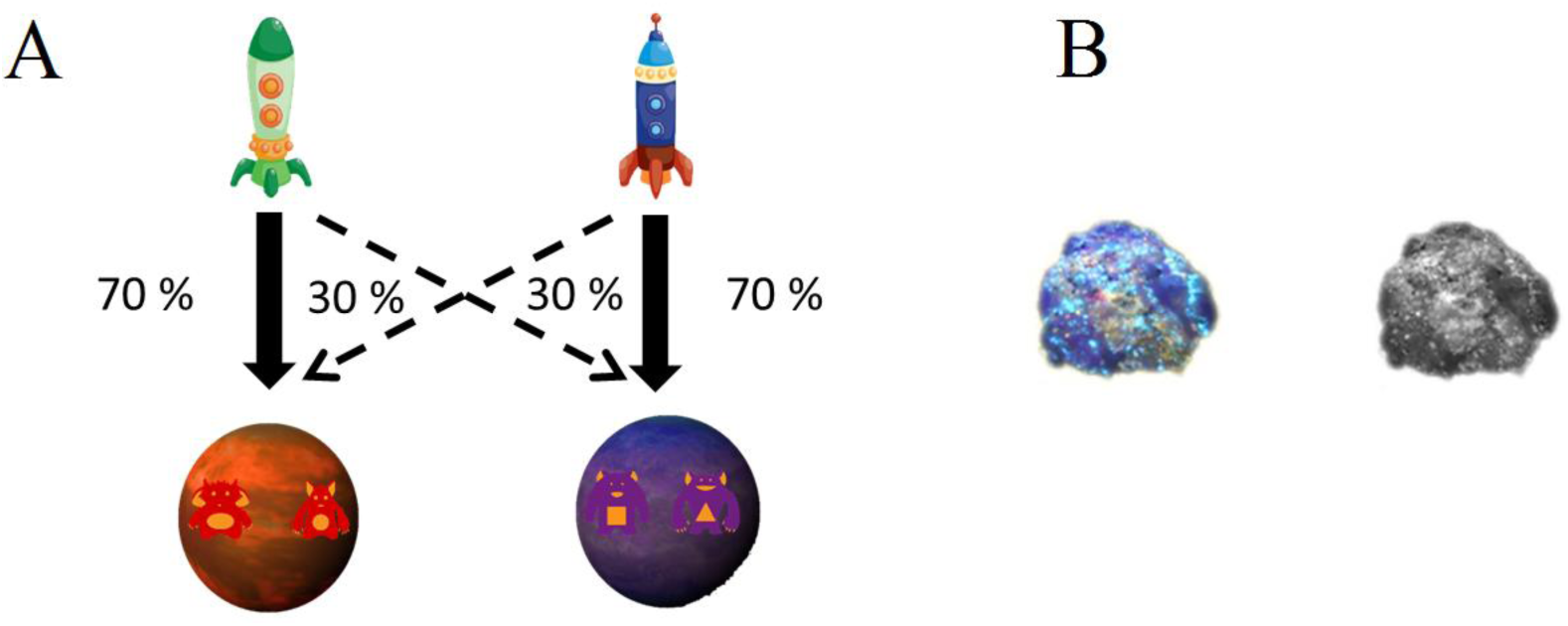
Transition structure and reward symbols. A. Following Daw et al. (2011) transition probability for each spaceship. Thick arrows indicate frequent transitions and thin arrows indicate infrequent transitions. In the Experiment, participants made two consecutive choices and were instructed to maximize their monetary earning by finding the pattern of choices that is more likely to result in a reward feedback to the second step choice. Participants chose between these spaceships on the first step and experienced this transition structure via observing which planet more frequently succeeds the choice. Note that the numbers beside each arrow indicates the probability of the corresponding transition. B. Treasure (colored) versus no treasure (grey) images. The treasure and no treasure serve as reward and no reward feedback to the second step choice. Here participants made a choice between two aliens that were differentially associated with reward and no reward.

Details of the trial events were as follows. On each trial, participants were first presented with an image of the home planet. After 300 ms, images of two spaceships appeared on the bottom left and bottom right corners of the screen. Participants selected the spaceships by pressing either “1” (for the spaceship on the left) or “0” (for the spaceship on the right) on a Qwerty keyboard. After a spaceship was selected, the image of the other spaceship was darkened (Figure 4c) and the selected spaceship moved slowly to the center of screen (600 ms, not shown). Then, the image of either the “red” planet or the “purple” planet (Figure 4d) was displayed at the center of the screen, indicating that the selected spaceship’s location of arrival on that trial. After 900 ms, two aliens associated with that planet—which were different in shape for the same planet and different in colour across planets—were shown on either side of the planet (Figure 4e). Participants selected the aliens by pressing the same keys (“1” for the alien on the left and “0” for the alien on the right). After one of the aliens was selected, the image of the other alien was darkened (Figure 4f) and the selected alien moved horizontally to the centre of the screen for 450 ms (Figure 4g). After 400 ms, one of the two images appeared on the screen and stayed on the screen for 600 ms, indicating whether participants won 5 cents (treasure image) versus 0 cents (no treasure image) (Figure 3b and Figure 4h).

**Figure 4.**
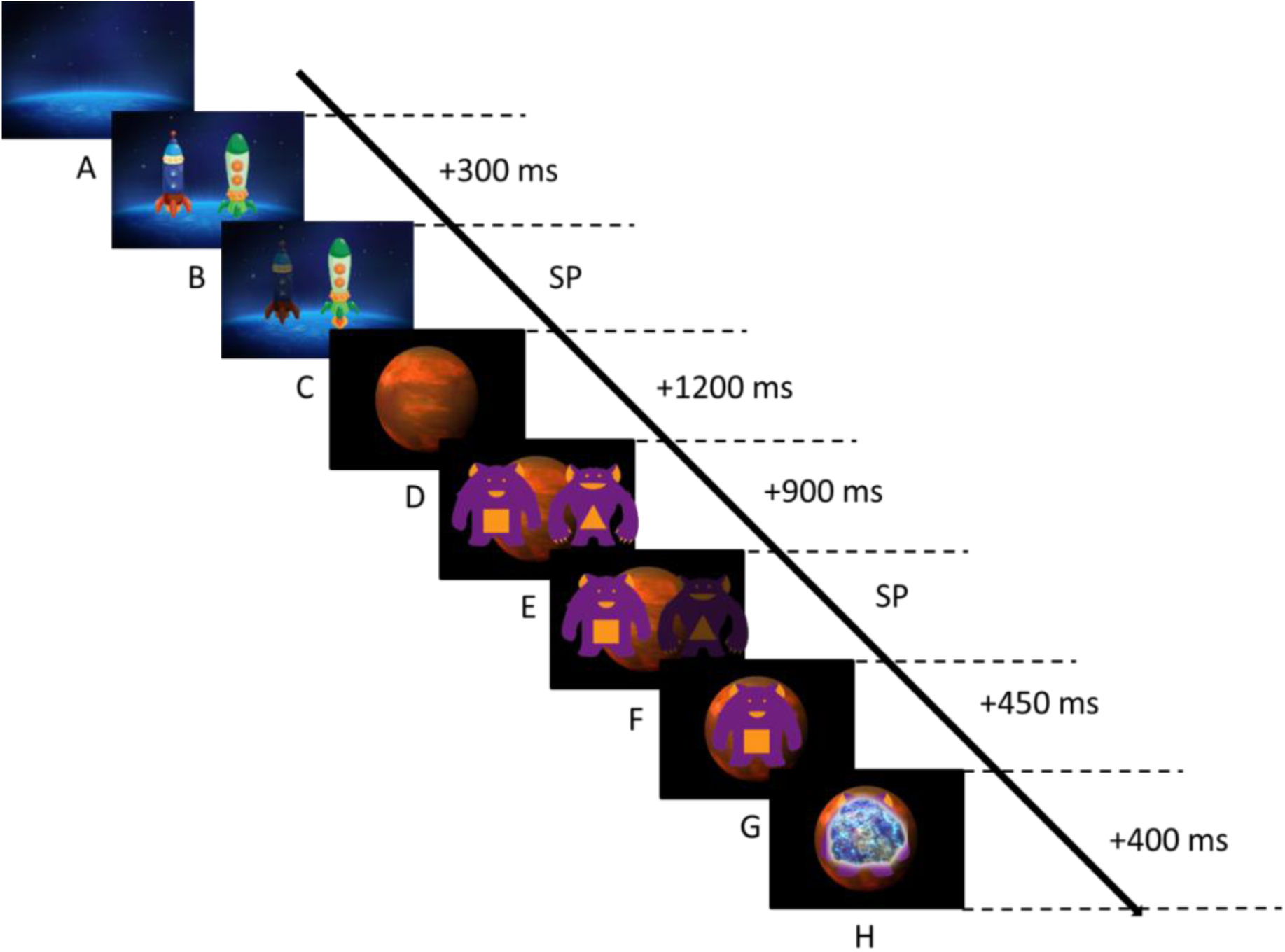
Task events for an example trial. The duration of each event is indicated in ms on the right side of each event itself; durations that depend on the participants’ pace are indicated as self-paced. A. An image of the home planet is presented. B. Two spaceship images are presented. C. Participants select one of the spaceships (self-paced), and the selected spaceship traverses to the center of the screen (not shown). D. A destination planet is shown on the screen. E. Two aliens appear on the planet. F. Participants select one of the aliens. G. The selected alien traverses to the centre of the screen. H. A feedback image appears indicating whether participants won 5 cents (coloured treasure) or 0 cents (grey rock) on that trial. Note that the distances between the images on each screen are not proportional to what was actually presented to the participants and are altered here for the purpose of illustration.

Similar to Daw et al. (2011), the payoff rate associated with each alien drifted probabilistically from trial to trial according to a random walk procedure; the specific random walk was chosen based on comparing the results of multiple random walk simulations and selecting the one in which the average expected probability of reward across all options was closest to 0.5 (Figure 5). All participants were exposed to the same action-reward contingencies.

**Figure 5.**
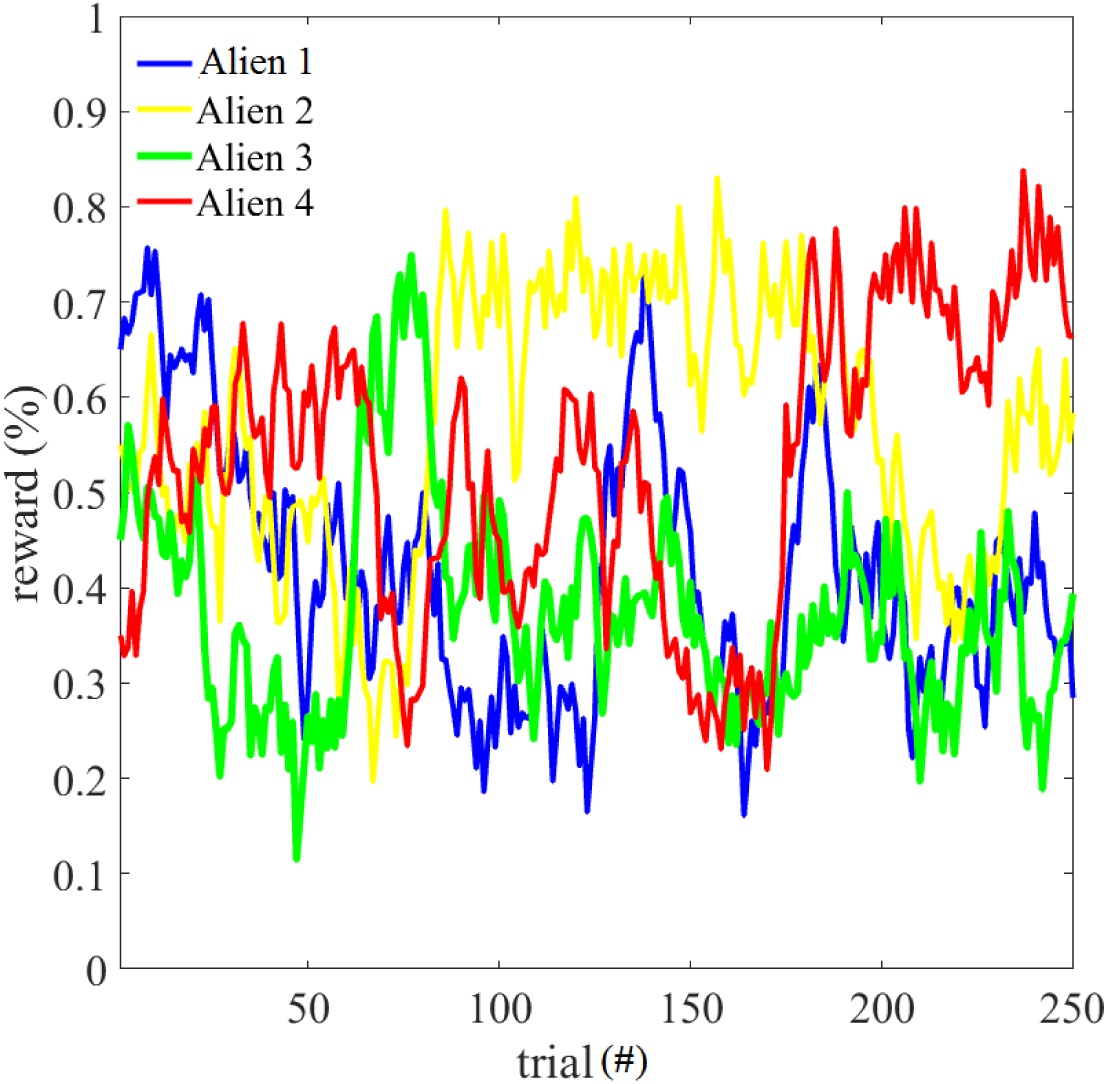
Probability that selecting each alien would yield reward on each trial across the entire experiment. The trial number is shown on x-axis and reward probability is shown on y-axis.

#### 3.1.3. Behavioral Data Analysis

To investigate whether participants’ first step choices were influenced by the transition type (frequent vs. infrequent) and outcome (reward vs. no reward) associated with the events in the preceding trial, we conducted a two-factor within-subjects ANOVA with factors outcome (2 levels: Reward, No-Reward) and transition (2 levels: frequent vs. infrequent) on the frequency of repeating the first step choice. We also conducted two planned comparisons: A repeated measures two-tailed t-test evaluating the effect of outcome on the frequency of stay when the transition event is a frequent (common) transition, and a repeated measures two-tailed t-test evaluating the effect of outcome on the frequency of stay when the transition event is infrequent (rare).

As has been suggested before (Daw et al., 2011), stay behavior (repeating the first step choice) on infrequent transitions provides a measure of the degree to which the decisions are based on model-based versus model-free systems. According to the model-based strategy, acquisition of reward on an infrequent transition should increase the likelihood of a switch on the upcoming trial, because the alternative first step choice is revalued based on the knowledge of transition probability. By contrast, following a model-free strategy, which is based on simple principles of reinforcement learning, agents are more likely to stay on their first choice when it is rewarded in the preceding trial regardless of the transition probability. Accordingly, recruitment of model-based and model-free systems exclusively should lead to choice behavior as depicted in Figure 6. More specifically, if the model-based system influences choices at a group level, then analysing frequency of stay on the first step choice should reveal an interaction effect between transition type and second choice feedback outcome in the preceding trial on the frequency of repeating the first step choice. To assess this effect, the tendency to use a lose-stay strategy for each individual was calculated in a manner similar to how group level interaction effect was calculated. If for each subject one calculates A, B, C, D as:

1. A is the frequency of repeating the same choice when the preceding trial involved a frequent transition and a reward
2. B is the frequency of repeating the same choice when the preceding trial involved a frequent transition and no reward
3. C is the frequency of repeating the same choice when the preceding trial involved a rare transition and a reward.
4. D is the frequency of repeating the same choice when the preceding trial involved a rare transition and no reward.

**Figure 6.**
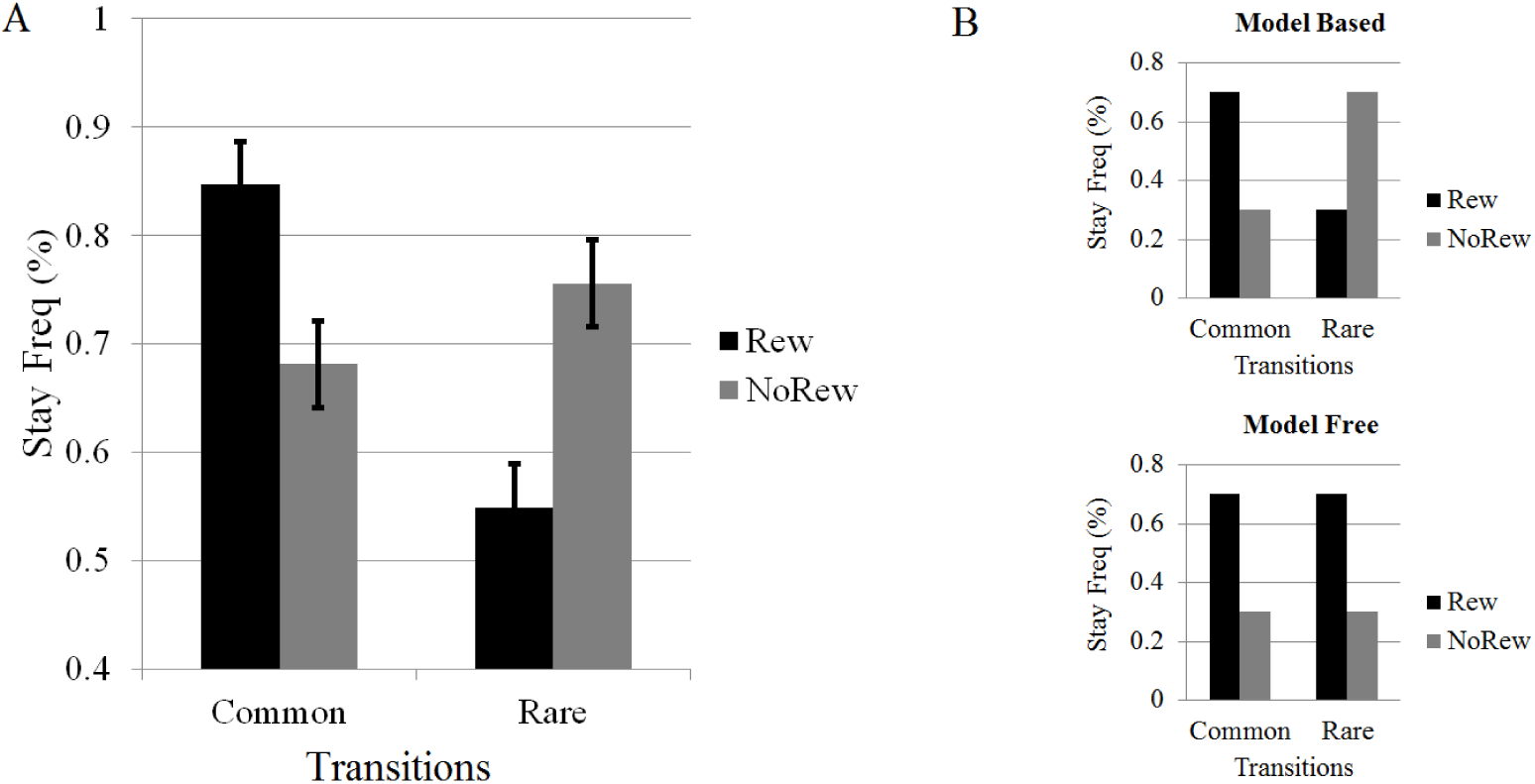
Stay frequency on the first choice depending on preceding trial’s outcome and transitions type. A. Participant stay frequency as a function of Transition (x-axis) and Outcome (Reward indicated in black and No-Reward indicated in grey). Error bars indicate 95% within subject confidence intervals (Masson, 2003). B. Schematic illustrating predictions for stay frequency according to model-based (top) and model-free (bottom) strategies, as a function of Transition and Outcome (note that the exact values are arbitrarily chosen; only the main effects are predicted by each model).

Then the effect of the transition event on the choice profile is reflected in the interaction term as (D-C) - (B-A) (see Friedel et al. 2014). A positive score on this measure indicates that the subject is likely to have a tendency to use a lose-stay strategy on the rare transition and in contrast use a lose-switch strategy on common transitions. Therefore, the pattern of choice behavior for subjects with positive score on this measure matches the pattern that can be predicted by model-based choice strategy. Hereafter, we will refer to the subject level effect of the transition event on the choice profile as the “individual model-based term”.

#### 3.1.4. EEG acquisition and preprocessing

The EEG data acquisition and preprocessing procedures were the same as in Experiment 1 except where explicitly stated.

#### 3.1.5. Time frequency analysis

All details of the time-frequency analysis are identical to Experiment 1 unless otherwise stated.

## a. Results

### 3.2.1 Behavior

A two-factor repeated measures ANOVA on stay frequency with transition type and outcome as factors revealed a significant main effect of transition type (F(1,39)=29.1, p < 0.05, MSE= 0.02) and an interaction between transition and outcome F(1,39)=34.4, p < 0.05, MSE= 0.04). However, there was no main effect of outcome (F(1,39)= 1.61, p > 0.05, MSE=0.01). The interaction effect shows that participants incorporated predicted transitions into their decision making. Indeed, a two-tailed repeated measures t-test revealed that participants were significantly more likely to repeat their choices on the first step if they were rewarded in trials with common transitions (*M*=0.16, *SD*=0.25, d = 0.64) compared to trials with rare transitions, *t*(39)=4.18, *p* < 0.05. In contrast, a two tailed repeated measures t-test revealed that on trials with rare transitions, participants were more likely to repeat their choices on the first step if they were not rewarded compared to if they were rewarded (*M*=0.21, *SD*=0.19, d=1.1), *t*(39)=6.59, *p* < 0.05). Note that for 33 out of 40 subjects the model-based influence term was positive, indicating that their decisions conformed to a model-based choice policy.

Further, reward receipt significantly modulated the choices at the second step: when the choice of the alien at the second step was rewarded, the subjects were more likely to choose that alien the subsequent time they arrived at the same planet (state) (M=0.87, which is significantly larger than chance level (0.5), t(39)=2.59, p < 0.05).

#### 3.2.2. Time-frequency response

Figure 7 shows the shows the grand-average difference in the time-frequency response to the transition stimuli in the 2–10 Hz frequency range. Contrary to our prediction, the mean difference in FMT power as a function of transition type was not significantly different from zero (M=1.95, SD=1.07), t(39)= 1.81, p > 0.05.

**Figure 7.**
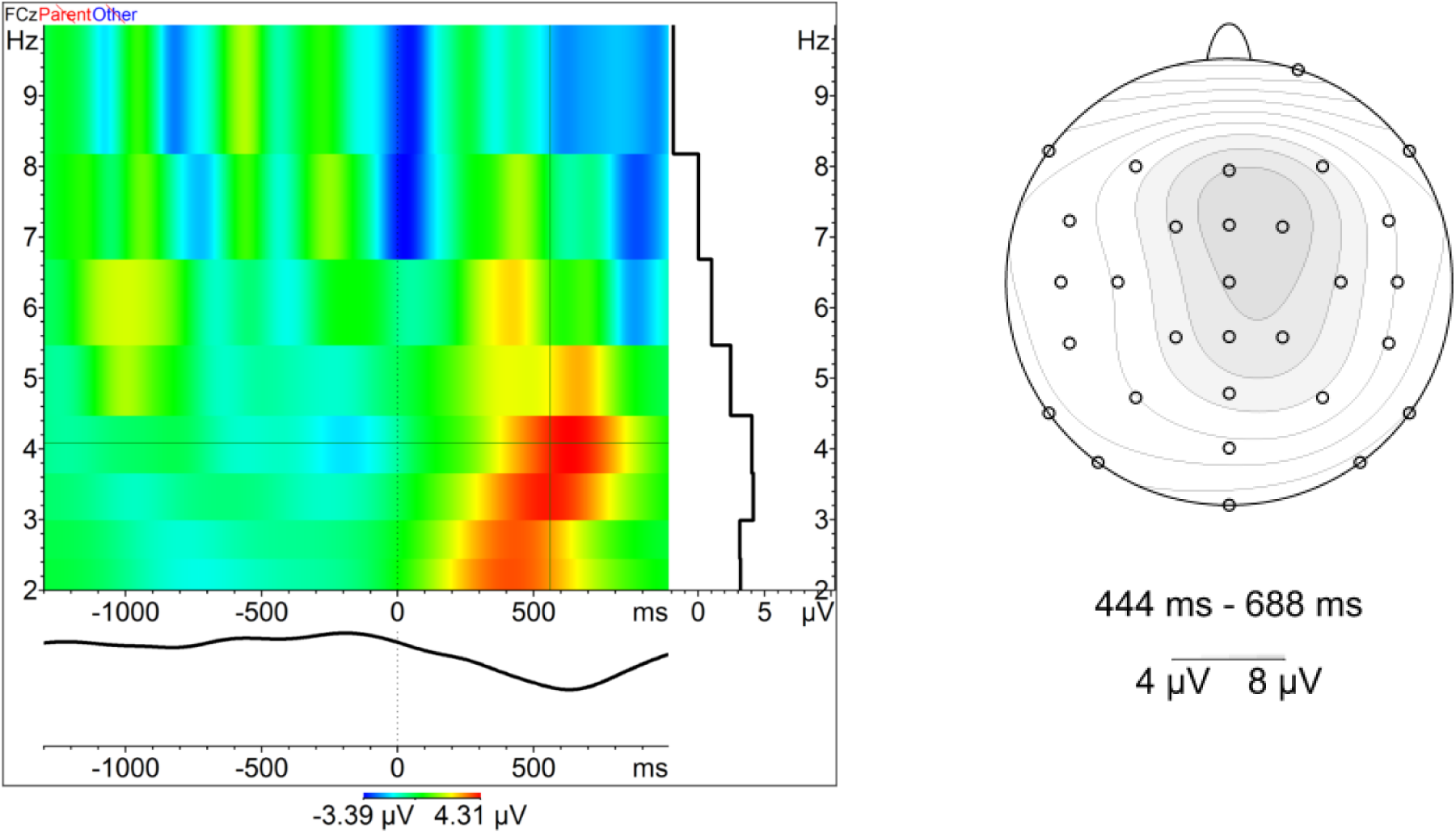
Top. The amplitude of difference in time-frequency response to rare versus common transition. The crosshairs denote the timing and the frequency of maximum power. Bottom. The scalp distribution associated with the average difference in time-frequency response in the 3–4 Hz range during the 444–688 ms interval post-stimulus.

However, inspection of Figure 7 suggested a later effect of transition type in the delta frequency range. An exploratory analysis revealed that the mean difference in delta power as a function of transition type in the 3–4 Hz frequency range and 444**–**688 ms interval post-stimulus confirmed that this difference was significantly different from zero (M=3.46, SD=4.87), t(39)=4.44, p < 0.05. This delta signal was characterized by a fronto-central scalp distribution (Fig 7, bottom). Importantly, the correlation between the model-based influence term and the difference in this delta signal to rare versus common transition stimuli was positive and significantly different from 0 (r(38)=0.348, p < 0.05), indicating relatively greater delta enhancement to rare transitions for participants who exhibited greater evidence of model-based behavior (Fig. 8). Lastly, visual inspection of the ERPs also revealed a difference in the 444**–**688 ms interval post-stimulus that was correlated with behavioral adjustments (see Supplementary online material).

**Figure 8.**
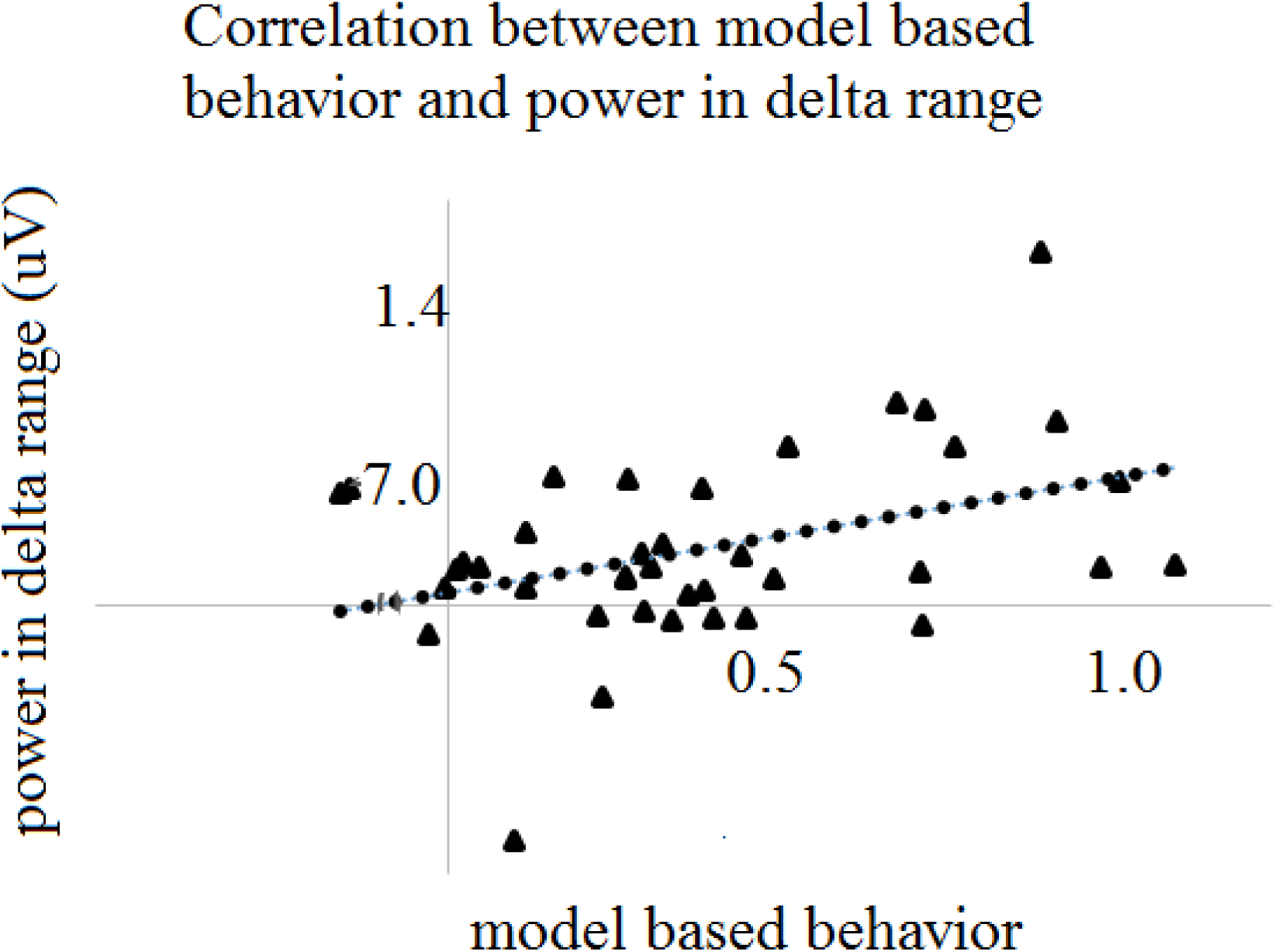
The correlation between model-based behavior with the difference in delta power to rare versus common transitions. The X-axis denotes individual model based behavior, computed as the differential frequency of repeating the same choice following rewarded versus unrewarded trials when the transition type is rare relative to the frequency of repeating the same choice following rewarded versus unrewarded trials when the transition type is common.

#### 3.2.3. Summary of the ERP findings

In accordance with Experiment 1, the amplitude of the P3 component was modulated by the transition event. However, in contrast to the Experiment 1, the transition event did not modulate the amplitude of the N2 above a detectable threshold. Importantly, the rare transitions increased the amplitude of the late positive complex (LPC) in the 444–688 ms interval.

### 3.3. Interim Discussion

Here we sought to replicate the findings from Experiment 1 – greater FMT to unexpected relative to expected state transitions -- in a reward-based planning task involving 2 sequential choices on each trial (Daw et al., 2011). Crucially, the choice on the first step is equivalent to the choice in the (one-step) task used in Experiment 1. Therefore, we expected to see a similar FMT response to the rare transitions relative to the common transitions. However, contrary to our predictions, there was no detectable FMT response to this event. In contrast, an exploratory analysis revealed that this transition event modulated delta oscillations in the period thereafter, and that this modulation exhibited a frontal central scalp distribution. Moreover, the difference in the delta response to the transition events (rare vs. common) was predictive of participants’ tendency to adopt a lose-stay strategy on the trials featuring a rare transition, which implicates the fronto-central delta response in model-based behavior.

## 4. General Discussion

The ability to predict the consequences of actions is crucial for successful planning. Learning about such contingencies relies on an model of the unlikely and likely outcomes of each action, i.e., the transition structure of the task. We reported the results of two experiments that instructed participants to pay attention to the likely and unlikely outcomes of each action in order to maximize their monetary earnings. Therefore, we expected that a neural mechanism responsible for processing SPEs would be recruited in both experiments.

In Experiment 1 we looked for neural evidence of SPEs in a task that did not involve reward feedback and therefore did not require explicit planning based on the task transition structure. To do so, participants were explicitly instructed to learn the likely outcomes of each action. Because the different states could not be associated with task-related valence, this process could not involve model-free learning (which relies on associating actions with future reward outcomes), and therefore elicited SPEs that were uncontaminated by RPEs. The pattern of choices on test trials revealed that participants were able to follow the action-outcome contingencies, confirming the development of a model-based transition structure. Concurrently, theta power was modulated in the predicted direction depending on whether the outcome was common or rare, and the scalp distribution of this effect was fronto-central, consistent with its identification as FMT. When evaluated in the context of prior literature indicating that FMT is generated in the aMCC (Cavanagh & Frank, 2014), these results indicate that aMCC responds to SPEs.

The purpose of Experiment 2 was to generalize these results to a task where subjects utilized the transition probabilities in order to plan for reward. However, in contrast to Experiment 1, because this experiment involved rewards, it allowed for a possible confound between SPEs and RPEs. Note that although the concurrent effects of RPEs and SPEs can be disentangled using advanced modeling approaches (see Sambrook, Hardwick, Wills, & Goslin, 2018), such approaches require the application of a linear regression on multiple variables (in the present case, SPE, RPE and the ongoing EEG), and so have relatively limited statistical power.

In Experiment 2, participants’ choices were affected by the transition type, indicating that they paid attention to the action-outcome contingencies. However, Experiment 2 did not replicate the FMT effect observed to state transitions in Experiment1. Several differences between the two experiments could account for this discrepancy. A first possibility, which we favour, is that the neural mechanism that updates the representation of the state-transition structure produces FMT only when learning is required. Therefore, Experiment 2 did not elicit FMT because the transition structure was explicitly instructed to the participants, whereas Experiment 1 did elicit FMT because it required participants to learn the transition structure by exposure to the task. In other words, the aMCC might produce SPEs when the transition model of the environment is being learned rather than applied (cf. Mas-Herrero & Marco-Pallarés, 2014).

Second, the task instructions in Experiment 2 emphasized the importance of the reward outcomes. This may have made reward outcomes more salient to the participants than the transition events, thereby attenuating the FMT response to such events. On this account, an alternative task design with task instructions that emphasized attention to the transition events should enhance the FMT response to those events. Other EEG phenomena have demonstrated similar sensitivity, for example, when feedback information is made salient by task instructions (Nieuwenhuis, Yeung, Holroyd, Schurger, & Cohen, 2004). Note that the subjects’ behavior clearly indicates that they attended to the transition events. Nevertheless, they may have paid less attention to these events in Experiment 2 than in Experiment 1.

Third, the planning component in Experiment 2 may have placed greater cognitive demands on decision-making and working memory processes relative to Experiment 1 (see Otto, Gershman, Markman, & Daw, 2013). In principle, this increased load could have interfered with the control mechanism producing the FMT response to rare transitions.

Finally, as described above, an RPE elicited by the transition event could have interfered with the production of the SPEs. In this case, the valence of the transition event in Experiment 2 would disrupt the production of FMT elicited by the transition event. For example, consider a participant who has learned the local action-outcome contingencies. On the first step of a trial, the participant could use this information to select the action that has a relatively high expected value on common transitions. Thus, a rare transition would lead to a state with lower expected value, yielding a negative RPE. For this reason, RPEs elicited by the transition events could confound the SPE analysis in Experiment 2.

Nevertheless, several considerations argue against this possibility. First, the rapidly evolving outcome contingencies are difficult to learn by design (Figure 5), so it is unlikely that the participants could generate stable predictions of the trial outcomes on the first step of each trial, nor that this would attenuate the FMT response to the event—especially given the large effect size observed in Experiment 1 (dz ∼ 3). This supposition is supported by the observation of a robust reward positivity ERP component elicited by the trial feedback (supplementary online materials), which is otherwise smaller in amplitude to predicted outcomes (Walsh & Anderson, 2012). Second, previous time-frequency studies have observed greater FMT power to negatively-valenced events (Cohen, Elger, & Ranganath, 2007; Cavanagh et al., 2012; HajiHosseini & Holroyd 2012); therefore, the negative RPE elicited by the rare transitions should elevate rather than cancel the effect of transition type on FMT power.

These considerations highlight the need for future studies to employ task paradigms that explicitly encourage attention to the transition events, in order to disambiguate the neural correlates of transition structure learning from those of other cognitive processes including decision-making and working memory functions.

–Further analyses indicated that this occurred during an interval when the late positive complex (LPC) ERP component showed a similar effect (SOM). Interestingly, the delta (and LPC) effect correlated with a tendency by participants to follow a model-based strategy as characterized by the Individual model-based term. This measure reflects how much participants incorporated the transition structure of the task in their choices. Given the significant correlation between this delta signal and the interaction term, the delta band activity appears to reflect participants’ reliance on a model-based decision making process. Notably, delta power has been hypothesized to code for motivationally relevant states (Knyazev, 2007). Therefore, we suggest that subjects who are more motivated to follow the task instructions (as revealed by elevated delta power) are more influenced behaviorally by the transition events. Interestingly, the medial prefrontal cortex has been said to be the source of waking delta oscillations, though this observation should be interpreted with caution given a scarcity of delta source localization studies (Harmony, 2013).

In addition to our study, 2 other experiments recently examined EEG correlates of the 2-step task. In particular, Sambrook, Hardwick, Wills, & Goslin, (2018) applied a modeling approach to the problem. Although they identified a cluster of neural activity at occipito-parietal channels that was correlated with SPEs, the size of this effect was very small. By contrast, Experiment 1 in our study eliminated the RPE/SPE confound in the two-step task by removing the reward element, which revealed a robust SPE. Similarly, Eppinger et al. (2017) reported an effect of transition type on the LPC that replicates our ERP findings reported in the SOM, though their effect was considerably smaller (dz=0.48) relative to that of our study (dz=1.17; supplementary online material). In summary, whereas our findings do not generally disagree with the previous results, ours is the first to report that SPEs elicit a robust FMT response and elevated delta levels that correlate with behavioral adjustments.

Note that sensitivity of a signal to the likelihood of events does not necessarily mean that the signal arises from a planning mechanism. In order to establish that FMT is sensitive to transition learning, future research should focus on predicting participants’ future choices based on trial-to-trial variations in signal size. In particular, if FMT power is predictive of participants’ choice behavior, then the signal must reflect subjects’ expectations about the action-outcome contingencies.

Taken together, the results of these two experiments indicate that these fronto-central oscillations—in the theta range in Experiment 1 and in delta range in Experiment 2—are responsive to transition probabilities. The results of Experiment 1 clearly implicate the aMCC in tracking action-outcome contingencies, which is in agreement with the results of previous studies. For example, Morris et al. (2017) showed that the aMCC tracks the causal association between an action and an upcoming state, and the PRO model advanced by Alexander and Brown (2011) holds that aMCC is responsible for detecting the non-occurrence of expected events. Likewise, other research has implicated the aMCC in constructing models of environmental contingencies. For example, O’Reilly et al. (2013) showed that aMCC’s BOLD level increases to surprising stimuli only when such stimuli call for a model update. Similarly, Karlsson, Terpo and Karpova (2012) showed that periods of behavioral uncertainty necessitating an update to the model are associated with resets of the ensemble activity of a network of neurons in the medial prefrontal cortex. These results are consistent with multiple observations that the aMCC reacts relatively strongly to surprising events (e.g., Ide, Shenoy, Angela, & Chiang-Shan, 2013; Mas-Herrero, Marco-Pallares, 2014, Payzan-LeNestour, Dunne, Bossaerts, & O’Doherty, 2013).

In previous work we developed a computational model that illustrates the role of aMCC in predicting environmental regularities (Shahnazian & Holroyd, 2018). Our present results support and extend this proposal by suggesting that the aMCC constructs an explicit model of environmental contingencies that can be applied in the service of planning, a process that is presumably carried out with a network of other brain areas including dorsolateral prefrontal cortex. When evaluated in the context of other evidence indicating a role for aMCC in representing actions hierarchically (e.g., Shahnazian, Shulver & Holroyd, in press; Holroyd & Yeung, 2012) and in effortful control (Vassena et al., 2018; Le Heron et al., in press), these results suggest that the aMCC may constitute a powerful neural mechanism for the adaptive control of behavior.

## Acknowledgment

We would like to thank Catherine Hartley for providing us with the experimental code for the two-steps choice task. This research was supported in part by funding from the Canada Research Chairs program and a Natural Sciences and Engineering Research Council of Canada Discovery Grant (312409–05) awarded to C.B.H

