## Supplementary material for "Electrophysiological correlates of state transition prediction errors"

**Event-related potentials (ERP)**

**Experiment 1.**

**ERP preprocessing and analysis**

#### EEG acquisition and preprocessing.

The EEG signal was acquired using a montage of 33 Ag/AgCl ring electrodes mounted on a nylon electrode cap according to the extended international 10-20 system (Jasper, 1958). Inter-electrode impedances were maintained below 20 kΩ using an abrasive conductive gel applied to each electrode. The electro-occulogram (EOG) was recorded for the purpose of ocular correction; horizontal EOG was recorded from the external canthi of both eyes, and vertical EOG was recorded from the sub-orbit of the right eye and electrode channel Fp2. Signals were amplified by differential amplifiers with a frequency response of DC 0.017-67.5 (90 dB per octave roll off) and digitized with a sampling rate of 250 per second. Digitized signals were stored on disk using Brain Vision Recorder software (Brain Products GmbH, Munich).

Epochs of 800 ms duration were created by extracting samples from 200 ms prior to 600 ms following the onset of the planet image from the continuous EEG, separately for each channel and subject. Data were baseline-corrected by subtracting the mean voltage during the 200 ms interval preceding planet image stimulus onset in each epoch from the post-planet stimulus voltages in that epoch. Muscular and other artifacts were removed using a ±100 microvolts threshold and a ±50 microvolts step threshold as rejection criteria.

To examine whether the ERPs averaged to the onset of the planet images were modulated by the spaceship-planet frequencies of association, we averaged separate ERPs for each block of trials according to whether that association was infrequent (rare) or frequent (common) on each block. We then examined the amplitude of the N2 and P3a ERP components to each condition in channel FCz. N2 amplitude was assessed using a base-to-peak measure by subtracting the mean voltage in the interval 176-224 ms post-stimulus from the mean voltage in the interval 232-280 ms post-stimulus recorded at channel FCz. P3a amplitude was calculated base-to-peak by subtracting the mean voltage in the interval 232-280 ms post-stimulus from the mean voltage in the interval 320-368. These time windows were identified by determining the peak of the P2 and N2 ERP components (N2) and the N2 and P3a ERP components (P3a), respectively, in the grand-average of the planet-locked ERPs at channel FCz. We focused on channel FCz based on the assumption that the activity in this channel reflects the activity of the aMCC. To examine scalp distributions, we calculated the difference wave by subtracting common transition related ERP from that of rare transition ERP.

ERP results. Figure S1 shows the time windows associated with P2, N2, P3 deflections. Note that the windows were created by identifying the P2, N2 and P3 peaks in the grand average ERPs to the **planet-image and then centering a 48 ms window for each component around the time of the associated peak.** Therefore, the windows were determined with an unbiased approach that is blind to the outcome likelihoods. Figure S2 shows the grand-average ERPs recorded at channel FCz to the planet images averaged across rare and common spaceship-planet transitions.

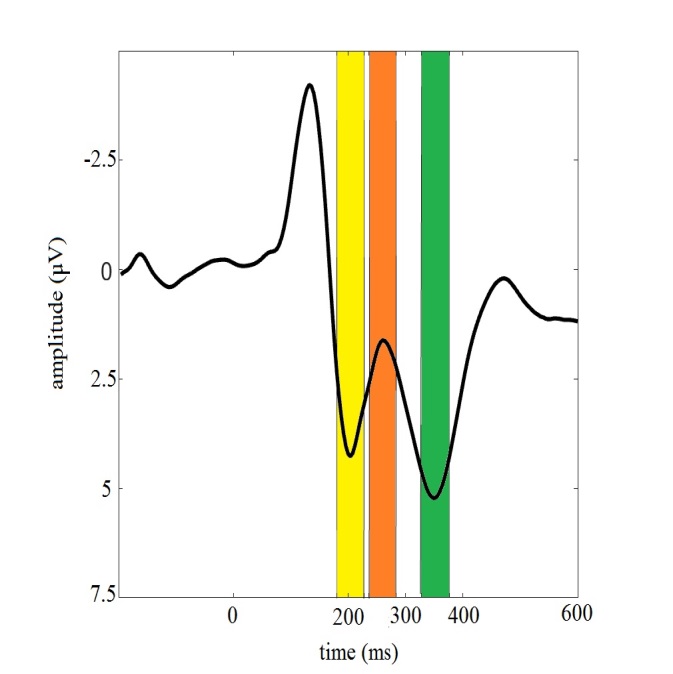

Figure S1. The latencies for P2, N2 and P3 peaks.

**The solid line depicts the planet-image locked ERP. The yellow stripe indicates the interval associated with the P2, the orange stripe indicates the interval associated with the N2, the green stripe indicates the interval associated with the P3.**

| 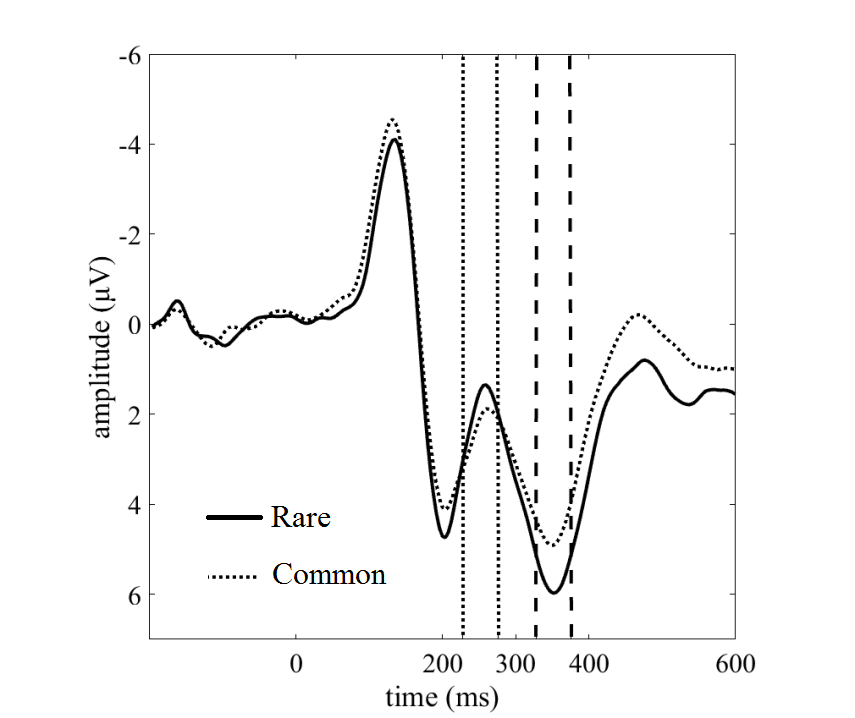 | 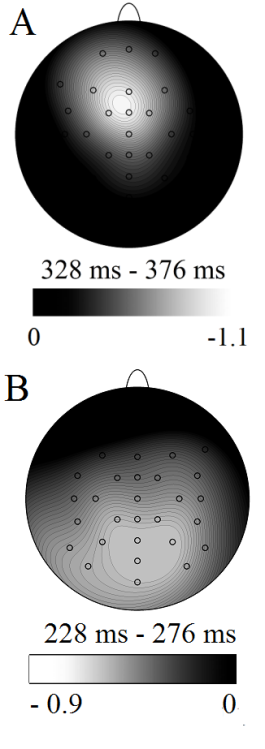 |
| --- | --- |

Figure S2. Grand-average ERP to rare and common transitions elicited by planet presentation, recorded at channel FCz.

Left. The solid line depicts the ERP to common transitions and the dotted line depicts the ERP to common transitions. The vertical dashed lines indicate the 328-376 ms interval post planet image presentation (i.e., the P3a window). Note that negative is plotted upward by convention. 0 ms in the x-axis is the planet image presentation onset. Right. A. The scalp distribution associated with the difference wave in the P3a interval as indicated by the vertical dashed lines in the left panel. Right. B. The scalp distribution associated with the difference wave in the N2 interval as indicated by the vertical dotted lines in the left panel. Note that the scalp distribution is not calculated based on the base-to-peak measure.

A two-tailed paired t-test analysis revealed a statistical difference in the amplitude of the N2 to infrequent versus frequent transitions (*M*=-0.87, $SD=1.48, d=0.58$), (*t*(19)=-2.65, *p* < 0.05). Note that the amplitude of the difference wave (not the base-to- peak measure) had a parietal distribution and was maximal at channel Pz (Figure S2). A comparable analysis also revealed a statistical difference for P3a amplitude, (*M=*1.58,$SD=2.06, d=0.76)$, *t*(19)=-3.42, *p* < 0.05.

**Experiment 2**

**ERP preprocessing and analysis**

The EEG data acquisition and preprocessing procedures were the same as in Experiment 1 except where explicitly stated. Separate ERPs were created for the frequent and infrequent transition events by averaging the trials separately for these conditions. N2 amplitude was assessed using a base-to-peak measure by subtracting the mean voltage in the 192-240 ms interval post-stimulus from the mean voltage in the 268-312 ms interval post-stimulus for the ERPs recorded at channel FCz. P3a amplitude was assessed using a base-to-peak measure by subtracting the mean voltage within the interval 268-312 ms post-stimulus onset from the mean voltage within 368-412 post-stimulus onset at channel FCz. Similar to Experiment 1, these time windows were identified by determining the peak of the P2 and N2 ERP components for the N2, and the N2 and P3a ERP components for the P3a, respectively, in the grand averages of the planet-locked ERP at channel FCz. P3 amplitude is often assessed in channel Pz, However, in our study, the P3 component as isolated with a difference wave approach exhibited a fronto-central scalp distribution, therefore we plotted the grand-average ERPs at channel FCz. Moreover, to examine how infrequent versus frequent transitions modulate the amplitude of ERPs to the events of interest, we conducted additional post-hoc analyses.

Additionally, we visually inspected the ERPs from -200 ms to 1000 ms post planet presentation to assess the qualitative differences in the grand average ERPs to frequent versus infrequent events recorded at channel FCz. In this examination, we observed that the infrequent transitions elicited a prolonged positivity in the 444-688 ms interval post-stimulus relative to the frequent transition (alternatively known as late positive complex or LPC see Bland and Schiffer, 2102). To assess whether this positivity was reliably expressed at the group level, we conducted an exploratory two-tailed repeated measures t-test on the mean voltage in the interval 444-688 ms on the planet-locked ERPs recorded at channel Pz (where the amplitude of difference wave in this interval was maximal; see Figure S5).

We also evaluated the amplitude of RewP (Sambrook & Goslin, 2015). We created ERPs locked to the treasure image (Reward ERP) and no-treasure image (No-Reward ERP). Reward positivity amplitude was calculated as the mean of the difference wave for Reward vs. No-Reward feedback in the interval 240-340 ms post-feedback onset, in accord with the prescription of a meta-analysis (Sambrook & Goslin, 2015).

To examine whether the different transition events modulate RewP amplitude, we conducted a two-factor within subject ANOVA with factors outcome (2 levels: Reward, No-Reward) and transition type (2 levels: frequent vs. infrequent) on RewP amplitude.

We also examined whether the amplitude of the transition-related ERPs were predictive of participants’ model-based behavior. To this end, we computed the correlation between the amplitude of difference wave for rare versus common transitions with the subject level strength of model based influence on stay-switch probabilities (see above). This analysis was done separately across subjects for each sample in the difference wave. We then identified the time-points for which this correlation value was significantly more than 0. Note that the analysis did not involve correcting for multiple comparisons.

#### ERP results.

Figure S3 shows the windows associated with P2, N2 and P3 deflections. Note that the windows are created by identifying the P2, N2 and P3 peaks in the grand average ERPs to the **planet-image,** and centering a 48 ms window for each component around the associated peak. Therefore, the windows are determined by an unbiased approach that was blind to the outcome likelihoods. Figure S4 shows the grand-average ERPs locked to the onset of the transition events. A two-tailed paired t-test on N2 amplitude did not reveal a statistical difference to the infrequent vs. frequent transitions, (M=0.05, $SD=1.45$ ) t(39)=0.2, p > 0.05. However, consistent with Experiment 1, a two-tailed paired t-test replicated the effect of transition on P3a amplitude, which was significantly larger to the rare transition relative to common transition (M=1.26, $SD=2.15, d=0.59$ ; t(39)=3.67, p < 0.05).

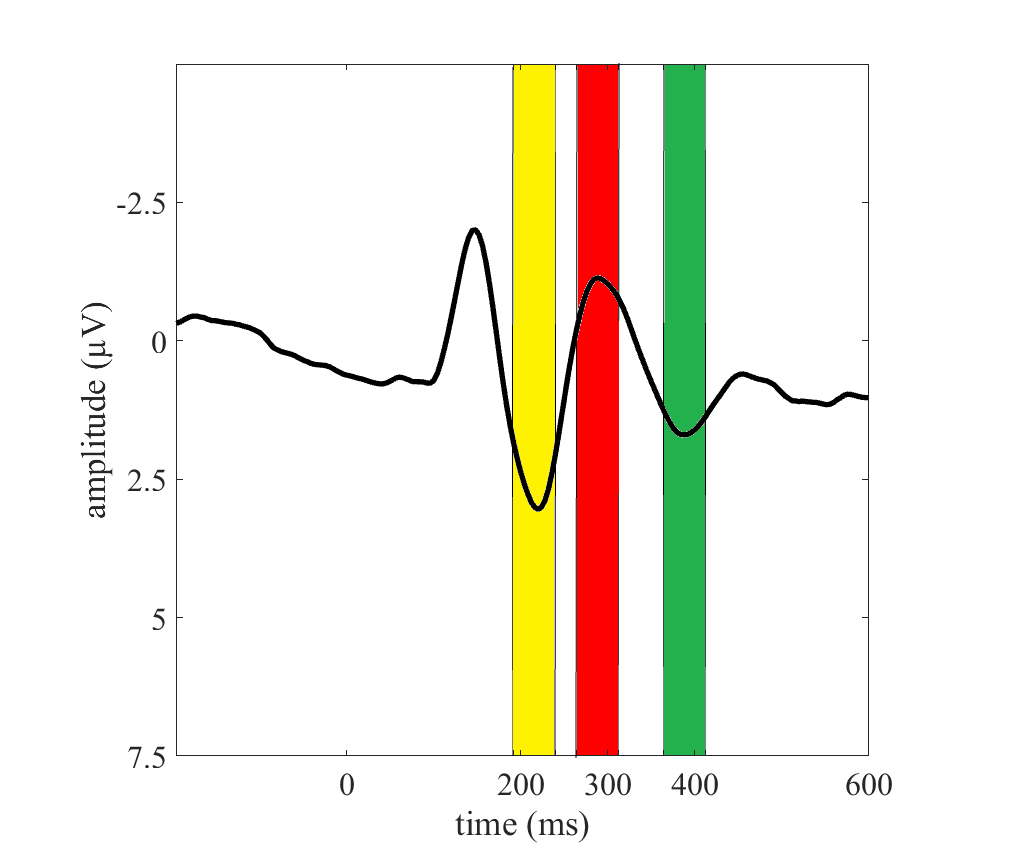

Figure S3. Time windows for the P2, N2 and P3a peaks.

The solid line depicts the planet-image locked ERP. The yellow stripe indicates the interval associated with P2, the red stripe indicates the interval associated with N2, the green stripe indicates the interval associated with P3a.

| 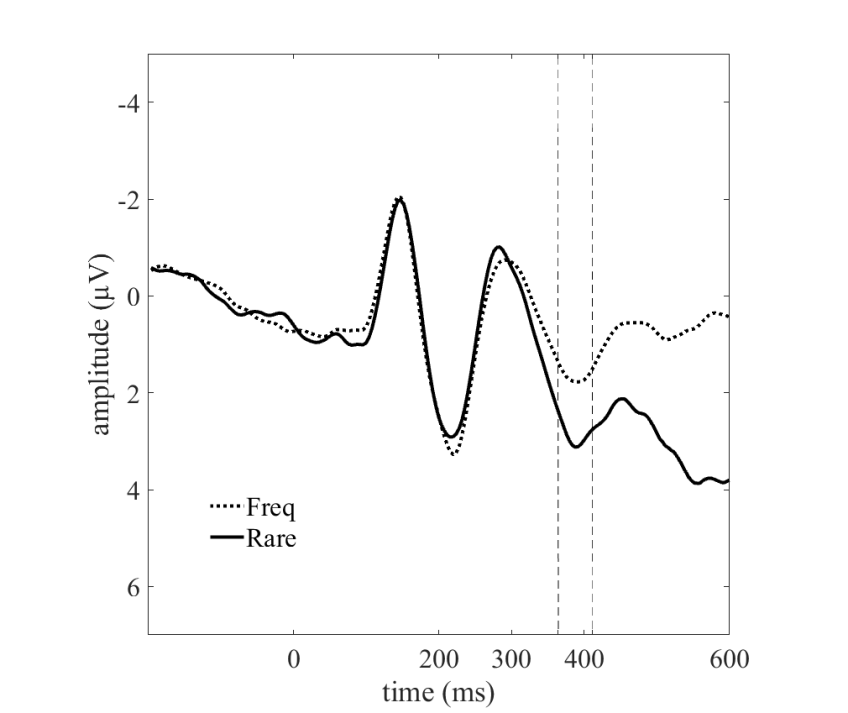 | 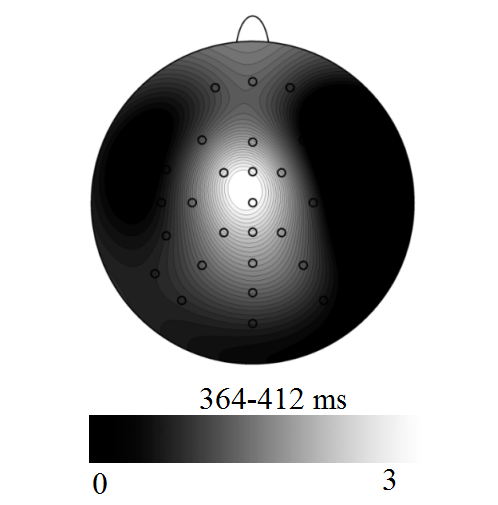 |
| --- | --- |

Figure S4. **Grand-average ERPs time-locked to** planet image onset recorded at channel FCz.

A. **The solid line indicates the ERP locked to the infrequent transition (Rare) and the dotted line indicates the ERP locked to frequent transition (Freq). 0 ms indicates planet image presentation onset. Vertical dashed lines indicate the RewP interval. B. The scalp distribution of the difference wave in the 368-412 ms interval post-stimulus. Lighter shading corresponds to more positive voltages.**

Additionally, visual inspection of grand-average ERPs recorded at channel FCz to the transition events suggested differential activity in a later period at about 444-688 ms post-planet onset (Figure S5).

| 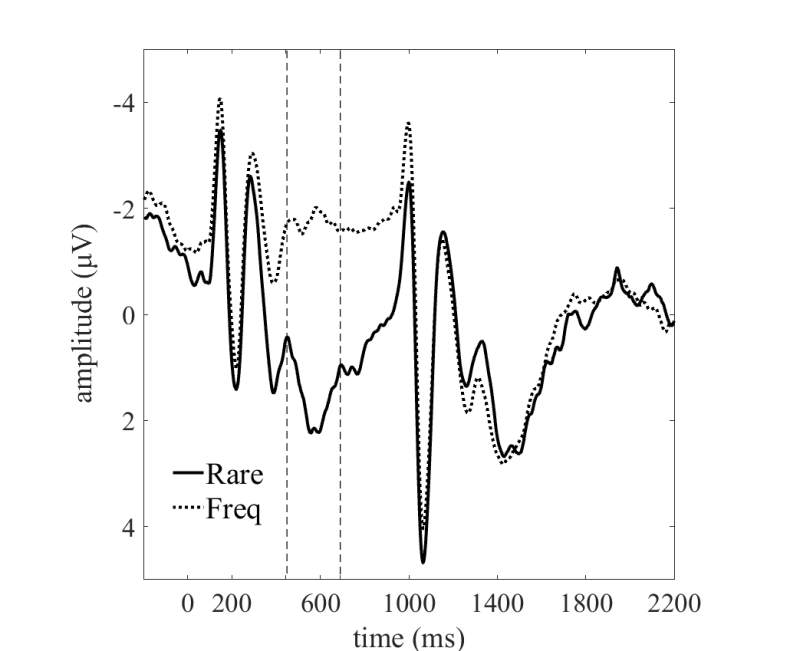 | 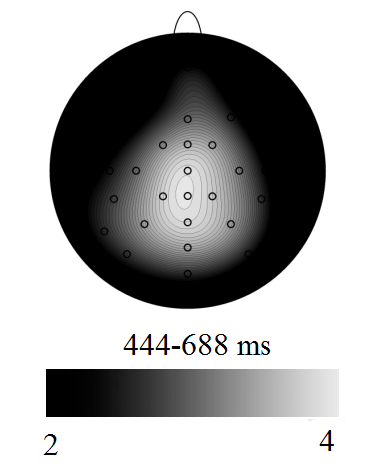 |
| --- | --- |

Figure S5. Prolonged positivity to rare transitions recorded at channel FCz.

A. **The solid line indicates the ERP locked to the infrequent transition (Rare) and the dotted line indicates the ERP locked to the frequent transition (Freq). 0 ms indicates planet image presentation onset. Vertical dashed lines indicate the interval 444-688 ms. B. The scalp distribution of the difference wave in 444-688 ms interval (indicated by vertical dashed lines) post-stimulus onset. Lighter shading indicates more positive values.**

A two-tailed repeated one sample t-test on the mean amplitude of the difference wave in the 444-688 ms interval post-stimulus onset revealed a significant effect of transition type, (*M*=3.29, *SD*=2.82, d=1.17), *t*(39)=7.37, *p* < 0.05, indicating that the waveform was more positive following rare transitions than following common transitions.

| 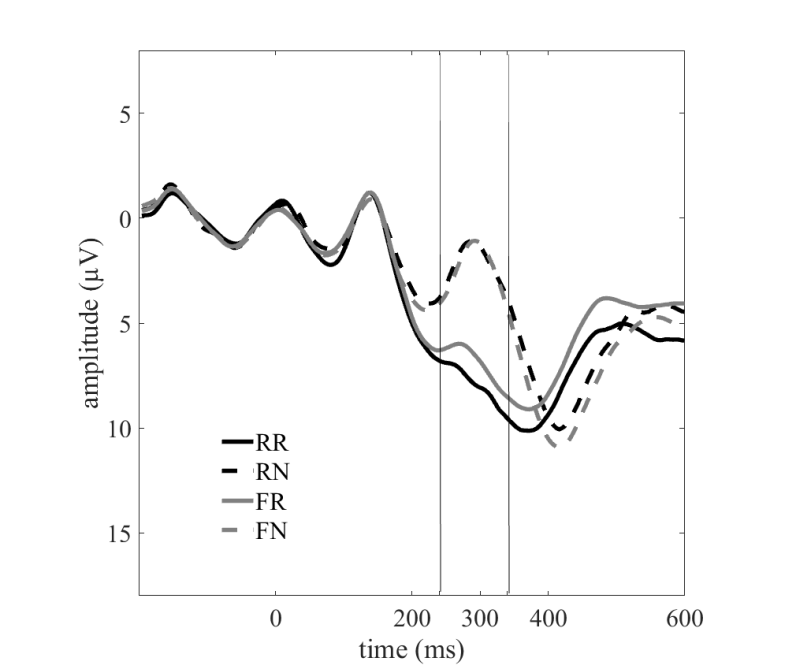 | 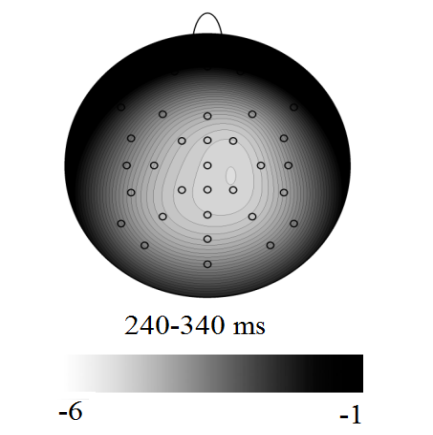 |
| --- | --- |

Figure S6. Feedback ERPs recorded at channel FCz showing the reward positivity (RewP).

RR denotes reward feedback following rare transitions. **RN denotes no-reward feedback following rare transitions. FR denotes reward feedback following frequent transitions. FN denotes no-reward feedback following frequent transitions. Dashed lines indicate no-reward feedback and solid line indicate reward feedback. Grey lines indicate a common transition type and black lines indicate a rare transition type. 0 ms indicates second step feedback presentation onset. The dotted vertical line indicates the RewP interval.** **B. The scalp distribution of the difference wave in the RewP interval for reward versus no-reward feedback ERPs. Lighter shading corresponds to more negative voltages.**

Figure S6 shows grand average ERPs to reward and no-reward outcomes conditioned on the preceding transition events. As seen by inspection, the scalp distribution of the difference wave created by subtracting the ERP to rewarded outcomes from the ERP to the unrewarded outcomes is centrally distributed and maximal at channel Cz, which is close to the fronto-central distribution typically reported in the literature (Walsh & Anderson, 2012). A two-factor repeated measures ANOVA with factors transition type and outcome on the amplitude of ERPs in the RewP interval following the feedback revealed a significant main effect of outcome (*F*(1,39)=213, p < 0.05 ,MSE= 4.82). However, there was no main effect of transition (*F*(1,39)= 1.7, p > 0.05, MSE=3.23) nor any interaction between transition and outcome (*F*(1,39)=3.44, p > 0.05 ,MSE= 3.35), indicating that RewP amplitude was not modulated by the transition events.

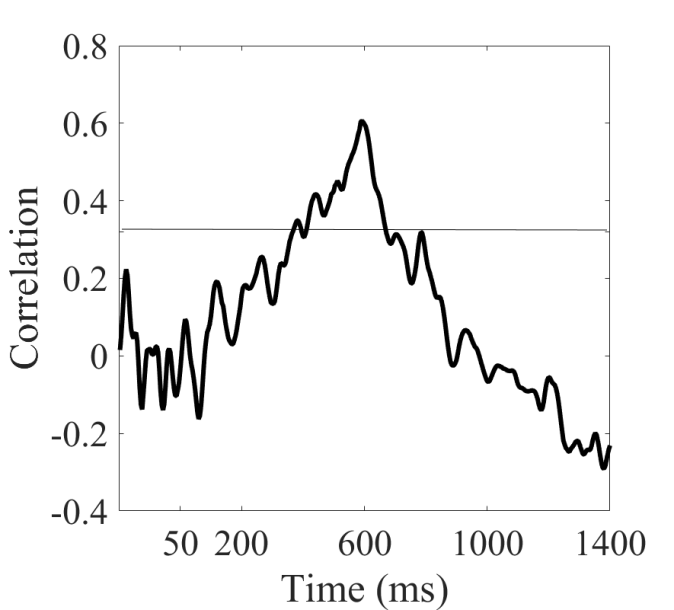

Figure S7. Correlation between the strength of model based influence and the amplitude of the difference wave for the different transition conditions.

The strength of model based influence assesses the influence of the model based system on choice behavior. The x-axis indicates time relative to the onset of the planet image and the y-axis indicates the correlation value. The solid line indicates the -critical value corresponding to the *p*-critical value = 0.05. **0 ms indicates planet image presentation onset**.

Figure S7 depicts the correlation between the subject-level tendency to use a lose-stay strategy on rare transitions and the difference wave ERP to the transition events for all of the time-points in the interval between -200 to 600 ms following the onset of the planet stimuli. Crucially, the correlation value exceeds the critical correlation value (*r*(38)$\boldsymbol{\approx}$0.31 for *p*-critical=0.05) in the interval 252-294 ms following planet stimulus presentation, which occurred during the late positive component (444-688) time window.
